# Pangenome genotyped structural variation improves molecular phenotype mapping in cattle

**DOI:** 10.1101/2023.06.21.545879

**Authors:** Alexander S. Leonard, Xena M. Mapel, Hubert Pausch

## Abstract

Expression and splicing quantitative trait loci (e/sQTL) are large contributors to phenotypic variability. Achieving sufficient statistical power for e/sQTL mapping requires large cohorts with both genotypes and molecular phenotypes, and so the genomic variation is often called from short read alignments which are unable to comprehensively resolve structural variation. Here we build a pangenome from 16 HiFi haplotype-resolved assemblies to identify small and structural variation and genotype them with PanGenie in 307 short read samples. We find high (>90%) concordance of PanGenie-genotyped and DeepVariant-called small variation, and confidently genotype close to 21M small and 43k structural variants in the larger population. We validate 85% of these structural variants (with MAF>0.1) directly with a subset of 25 short read samples that also have medium coverage HiFi reads. We then conduct e/sQTL mapping with this comprehensive variant set in a subset of 117 cattle that have testis transcriptome data and find 92 structural variants as causal candidates for eQTL and 73 for sQTL. We find that roughly half of top associated structural variants affecting expression or splicing are transposable elements, such as SV-eQTLs for *STN1* and *MYH7* and SV-sQTLs for *CEP89* and *ASAH2*. Extensive linkage disequilibrium between small and structural variation results in only 28 additional eQTL and 17 sQTL discovered when including SVs, although many top associated SVs are compelling candidates.

## Introduction

Assigning functional information to genetic variants is challenging. Genome-wide association studies (GWAS) have revealed many QTL in cattle (Freebern et al. 2020; Fang and Pausch 2019), and other species (Yengo et al. 2022; Filiault and Maloof 2012), but require substantial *a priori* knowledge of phenotypes or traits of interest. Alternatively, expression quantitative trait loci (eQTL) mapping can use “molecular phenotypes”, such as RNA abundance, to identify regulatory variants, which may contribute to inherited trait variation (e.g., carcass yield (Wang et al. 2022; Leal-Gutiérrez et al. 2020), male fertility (Mapel et al. Companion paper), and female fertility (Forutan et al. 2023)). Similarly, variants that are associated with alternative splicing or differential isoform usage can be identified through splicing QTL (sQTL) mapping (e.g., milk production (Xiang et al. 2018) and male fertility (Mapel et al. Companion paper)). In particular, sQTL have been suggested as a leading candidate for explaining a substantial portion of complex trait and disease heritability (Xiang et al. 2022). Alternative splicing can also affect gene expression and associated variants may also appear as eQTL (Yamaguchi et al. 2022).

Detecting e/sQTL relies on both accurate and complete quantification of RNA abundance as well as availability of matched genotypes from the same samples. Recently, several long read cohorts have demonstrated the importance of including structural variants (SVs) in explaining phenotypic variation in human (Beyter et al. 2021), tomato (Alonge et al. 2020), and rice (Shang et al. 2022). However, most e/sQTL studies, particularly those in livestock, primarily rely on short read sequencing (Littlejohn et al. 2016) or genotyping arrays (Cai et al. 2023; Liu et al. 2020) to assess genomic variants in enough samples to ensure sufficient statistical power to detect associations with molecular phenotypes. SVs, such as indels larger than 50 bp, have thus been predominantly neglected in GWAS and e/sQTL studies, despite contributing substantially to phenotype variation (Alonge et al. 2020; Scott et al. 2021). Some recent work has used short reads from various cattle breeds to call SVs (Zhou et al. 2022a; Lee et al. 2023; Bhati et al. 2023), but are primarily restricted to deletions and duplications and require extreme filtering to remove false positives. Long and accurate sequencing reads, like PacBio HiFi and those produced with ONT r10 chemistries, have the potential to access both small (including SNPs and indels smaller than 50 bp) and structural variants, but are costly when sequencing entire mapping cohorts.

A recent intermediate approach, PanGenie (Ebler et al. 2022), produces a “pangenome” variation panel from high-quality, haplotype-resolved assemblies (which can access all scales of variation). Additional samples can then be efficiently genotyped against this panel using *k*-mers. Crucially, short read sequencing can be used to produce these *k*-mers, enabling genotyping of both small and structural variants in existing biobank-sized short read cohorts. Earlier pangenome genotyping methods (Chen et al. 2019; Sirén et al. 2021) relied on more laborious sequence alignment to genotype samples, impeding scaling to larger cohorts.

Here, we create a pangenome variation panel from 16 haplotype-resolved cattle assemblies: six Original Braunvieh (OBV), nine Brown Swiss (BSW), and one Piedmontese (PIE). Subsequently, we use PanGenie and DeepVariant on 307 short read sequence samples of predominant BSW and OBV ancestry to generate a comprehensive catalogue of small and structural variation. We conduct e/sQTL analyses on 117 deeply sequenced total RNA samples from testis tissue, and reveal QTL missed by the short read-only dataset as well as QTL where SVs are the causal candidate. While we show PanGenie cannot capture rare SVs, potentially missing large effect size QTL, we demonstrate this approach does improve e/sQTL mapping over using only short reads.

## Results

We created the pangenome variant panel using 16 HiFi-based haplotype-resolved cattle assemblies, including four previously published assemblies (two Original Braunvieh, one Brown Swiss, and one Piedmontese; (Crysnanto et al. 2021; Leonard et al. 2022)), eight newly generated assemblies from previously published data (four Original Braunvieh and four Brown Swiss; (Leonard et al. 2023)), and four assemblies from new data (four Brown Swiss). All assemblies were aligned to the cattle reference genome, ARS-UCD1.2 (Rosen et al. 2020), followed by calling variants from the alignments in confident regions (as described by PanGenie). Through this approach, we identified 12,918,792 SNPs, 3,123,739 indels shorter than 50 bp, and 53,297 structural variants longer than or equal to 50 bp. The three OBV individuals (six haplotypes) were a sire-dam-offspring trio, allowing us to estimate the SNP and SV mendelian inconsistency rate in the pangenome as 1.06% and 2.32% respectively.

We checked this set of assembly-derived SVs against SVs called directly from the HiFi reads with Sniffles and confirmed a high level of overlap (86%) as expected (Figure 1A). However, there was some disagreement, particularly for large insertions exceeding the average HiFi read size (Figure 1B). In these circumstances, the read alignments end in soft or hard clips on both ends of the insertion, and the SV cannot be directly detected (Supplementary Figure 1). Since the assemblies are effectively a single read with megabase-scale length, they can cleanly resolve larger insertion SVs. Deletions larger than the read length can generally still be directly detected. There were spikes of SV frequency for both insertions and deletions approximately of size 1.3 Kb, largely confirmed by RepeatMasker to be endogenous retrovirus (ERV) sequence.

**Figure 1.**
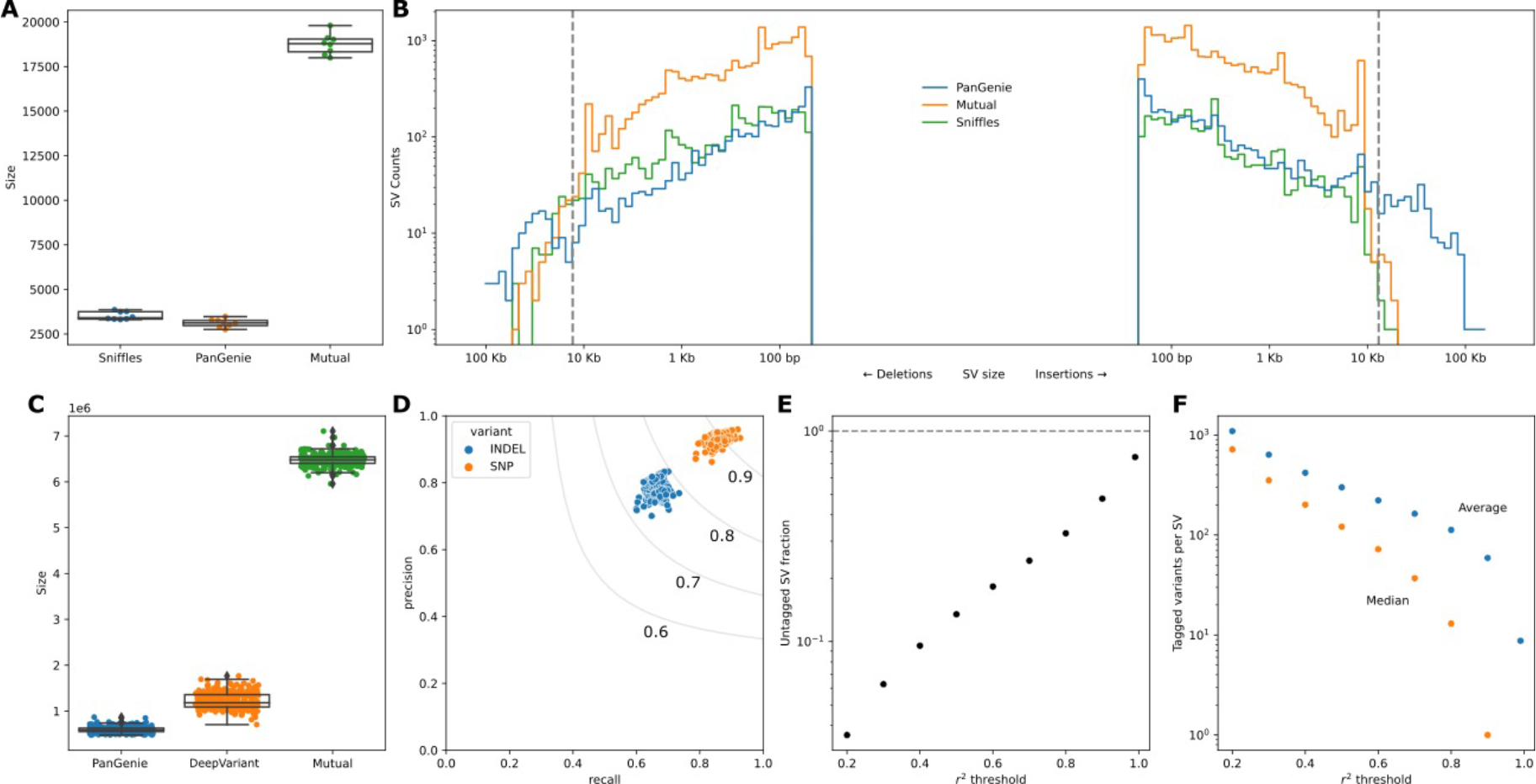
Concordance of variants genotyped by PanGenie. (A) SV overlap between PanGenie and Sniffles for the 8 individuals used to create the pangenome variant panel. (B) SV size distribution for the groups in (A). The grey dashed lines indicate 15 Kb, the average read length for the HiFi reads used by Sniffles. (C) Small variant overlap between PanGenie-genotyped variants and DeepVariant-called variants for the 307 short read samples. (D) Precision and recall for the 307 samples from (C). The grey lines are the F-score boundaries for the indicated values. (E) Fraction of all SVs tagged by small variants at different thresholds of r^2^ within a linkage window of 1000 Kb across the 307 samples. (F) Average and median number of variants that tag each SV across different r^2^ thresholds.

With PanGenie, we genotyped all the pangenome variation for 307 Braunvieh samples (consisting of Brown Swiss/Original Braunvieh/mixed breeds originating from a common ancestral population) using short sequencing reads. We further supplemented this genotyped variation by directly calling variants with DeepVariant on the 307 samples and merged the PanGenie-called and DeepVariant-called variation into a “PanGenie+” set. The larger sample size for DeepVariant (307 samples versus 8 individuals with haplotype-resolved assemblies) meant more small variants were called, although the majority were mutually present (Figure 1C). The overwhelming majority of SNPs and SVs in the pangenome variation panel were present in the larger cohort, 98.8% and 96.2% respectively, while small indels (<50 bp) were more frequently missing (80.8% present). The genotype concordance of the calls was also high, with mean F-scores of 0.90 and 0.72 for SNPs and indels respectively across the 307 samples (Figure 1D). There were also four distinct sire-dam-offspring trios in the 307 samples, which we used to validate the genotyping accuracy. The mendelian inconsistency rate was 1.15% and 4.46% for SNPs and SVs respectively. We also confirmed the genotyped small and structural variants were both independently able to recover the expected population structure through principal component analyses, although the more complete small variant set explained slightly more of the structure (Supplementary Figure 2).

We also examined the linkage disequilibrium (LD) between small variants and SVs, finding that nearly 70% of SVs are strongly tagged (r^2^>0.8) by small variants within a 1 Mb cis-window, while only approximately 5% of SVs are poorly tagged (r^2^<0.2) (Figure 1E). SVs were tagged by an average of 116 variants (median of 15 variants) within that window above the strongly tagged threshold (Figure 1F).

### Variant discovery in cohort subset with long reads

We also collected moderate coverage (12.9±1.4-fold) of PacBio HiFi reads on 25 samples of predominant Braunvieh ancestry and called structural variants from the long read alignments using Sniffles. We find that even a small number of samples captures a large portion of SVs present in a given population (Figure 2A), and we estimate that roughly 100 samples would likely capture nearly all SVs that segregate in a typical taurine cattle breed such as Braunvieh, finding only approximately 100 new SVs per additional sample beyond this population size (Supplementary Figure 3). Crucially, 69% of SVs discovered through the 25 long read alignments were already present in the PanGenie variant set and genotyped into the larger population (Figure 2B), rising to 85% when considering only SVs with allele frequency greater than 10% (Figure 2C). However, there were also 15,930 SVs that were not in the PanGenie variant set. As such, there is a non-negligible portion of SVs that could only be discovered through including additional assemblies into the PanGenie variant set or directly calling SVs with long reads on each sample in the e/sQTL set.

**Figure 2.**
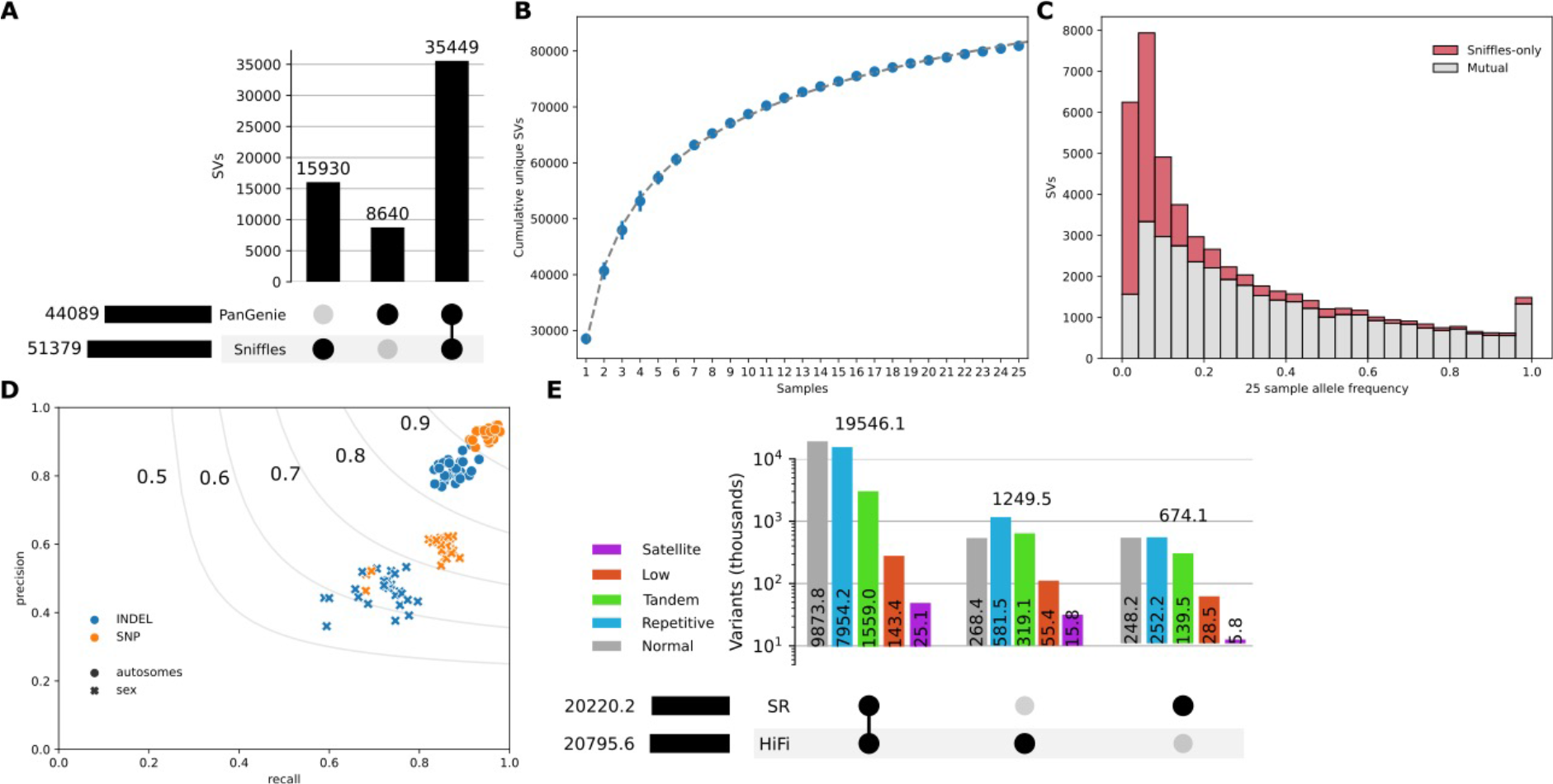
Comparison of variant calling with a small long read cohort. (A) SV intersection between PanGenie (called from 8 individuals with haplotype-resolved assemblies) and Sniffles (called from 25 HiFi read samples). (B) SV saturation for 25 HiFi read samples. Markers indicate the mean value of unique SVs over 10 random shuffles of sample order, and error bars represent standard deviation. The dotted line is a fitted curve of the form *f(x) = ax^-b^ + c*, predicting saturation at approximately 175,000 SVs. (C) SV overlap for different allele frequency (based on the 25 samples) bins. (D) Small variant accuracy of HiFi-based and short read-based calls, taking the short read data as truth, stratified by autosomes and sex chromosomes for SNPs and indels. Large markers indicate the mean over the 25 samples. (E) Small variant intersections between HiFi-based and short read-based calls in genomic regions identified as centromeric satellites, low mappability, tandem repeats, repetitive, and “normal” (all other regions). A large proportion of variants called in the challenging regions were unique to HiFi-based alignment and calling.

We were also able to compare small variant accuracy between HiFi and short reads in the 25 samples with approximately 10-fold coverage of both sequencing approaches. Notably, while there are minor differences for autosome-wide alignments between HiFi and short reads, with HiFi read alignments covering only 0.3% more of the autosomal bases than the short read alignments, there is moderate and large effect for the X and Y chromosomes respectively: 3.5% and 31.5%. The improved alignments in the sex chromosomes contributed most of the additional variants called by HiFi reads over short reads (Supplementary Figure 4). Taking the short read variants as truth, the mean SNP and indel F-score was 0.92 and 0.82 respectively (Figure 2D), where the higher recall than precision is largely due to the additional variants called by HiFi reads. Similarly, we observed that HiFi-based alignments (at comparable coverage) called substantially more variants in regions annotated as centromeric satellites, low mappability, tandem repeats, and repetitive, resulting from inconsistent and lower-quality short read alignments, while the number of variants in “normal” regions was comparable (Figure 2E).

### cis-expression QTL mapping

After splitting multiallelic variants and filtering at 1% minor allele frequency in the PanGenie+ set, 20,931,316 variants remained for downstream analyses, including 17,439,736 SNPs, 3,449,049 small indels, and 42,531 SVs (Table 1). There were 8,355 SVs larger than 1 Kb of which 2,103 were larger than 5 Kb. Since the SVs were only genotyped through PanGenie while small variants were also called directly, there were fewer rare SVs filtered out compared to SNPs (Supplementary Figure 5).

**Table 1.** Breakdown of variants for SNPs, small insertions and deletions (<50 bp), and SV insertion and deletions (≥50 bp) for the total merged PanGenie+ variant set and the MAF filtered variant set.

|  | SNPs | Small<br>insertion | Small<br>deletion | SV<br>insertion | SV<br>deletion |
| --- | --- | --- | --- | --- | --- |
| PanGenie+ | 23,278,277 | 2,954,293 | 2,502,528 | 28,087 | 21,033 |
| PanGenie+ MAF ≥ 0.01 | 17,439,736 | 1,852,334 | 1,596,065 | 24,193 | 18,338 |

We then investigated the impact of SVs on gene expression in a subset of 117 mature bulls for which we also had deep total RNA sequencing from testis tissue, with 257±35 million paired-end reads per sample. After aligning to the cattle reference genome and annotation (Ensembl release 104), followed by quantifying expression as transcripts per million (TPM), we retained 19,440 genes for cis-expression QTL (cis-eQTL) mapping. We ran a permutation analysis to determine the significance thresholds, followed by a conditional analysis, finding 3,677,218 associated variants for 15,406 eQTL (11,030 expressed genes [eGenes]). Of those variants, 6,985 were SVs (including 1,412 and 97 SVs longer than 1 and 10 Kb respectively). Association testing in a relatively small cohort of animals with widespread linkage disequilibrium often produces identical test statistics for multiple nearby variants. As the most significantly associated variant is not necessarily the causative variant, we also considered variants with conditional significance within 1.5x of the top variant (adapted from (Sanchez et al. 2017)) as candidate causal variants. We find 92 SV-eQTL, where 25 have eSVs as the unique-top variant (Figure 2A) and 58 eQTL where the top variant is an SV that is in near-perfect linkage disequilibrium with a small variant (Figure 2B).

We also performed the permutation and conditional analyses using only the DeepVariant variant set, representing the confident set of variation accessible with only short reads. There were 3,613,475 variants associated with the expression of 11,061 eGenes. All eGenes found uniquely with the short read dataset were just missed by the significance threshold in the full dataset, suggesting they are of marginal importance (Figure 2C). On the other hand, there were 29 eGenes found only with the PanGenie+ dataset, including 4 where the top eVariant was an SV (and the remaining were typically small indels within tandem repeats).

We further examined in more detail several eGenes that are affected by SVs identified with the PanGenie+ set (Supplementary Table 1). For example, we identified a strong cis-eQTL approximately 14 Kb downstream of the annotated translation termination codon of *STN1* (*ENSBTAG00000015019*) encoding STN1 subunit of the CST complex (Figure 4A). This cis-eQTL was significantly associated with 672 variants, although one of the top variants (p=1.99e-22) was a 5.9 Kb deletion containing 3.9 Kb of DNA transposons, RTEs, and ERV-LTR elements, occurring with a frequency of 32% in the 117 animals. The deletion is in high LD (r^2^=0.923) with the top SNP (Chr26:24452023 bp) and the association signal only slightly lower. *STN1* is moderately expressed (13.59 ± 1.92 TPM) in testis but the deletion reduces STN1 mRNA abundance (effect size [ß] of −1.11). Closer inspection of the eQTL also revealed limitations of the current functional annotation of the bovine reference genome. The Ensembl annotation of STN1 contains five transcripts whereas the Refseq annotation suggests ten isoforms, of which eight are expressed in testis including one (XM_024985601.1) that has an intron overlapping the deletion (Supplementary Figure 6A). While the deletion reduces the expression of three isoforms including the canonical isoform (NM_001076849.1), it increases the abundance of five other isoforms (Supplementary Figure 6B). Since the canonical isoform is more abundant than all other isoforms, the deletion overall reduces STN1 expression. We corroborated that the SV-eQTL not only impacts the overall expression of STN1 but also its relative isoform abundance, as this SV is also strongly associated with a splicing QTL (p=3.96e-22).

**Figure 3.**
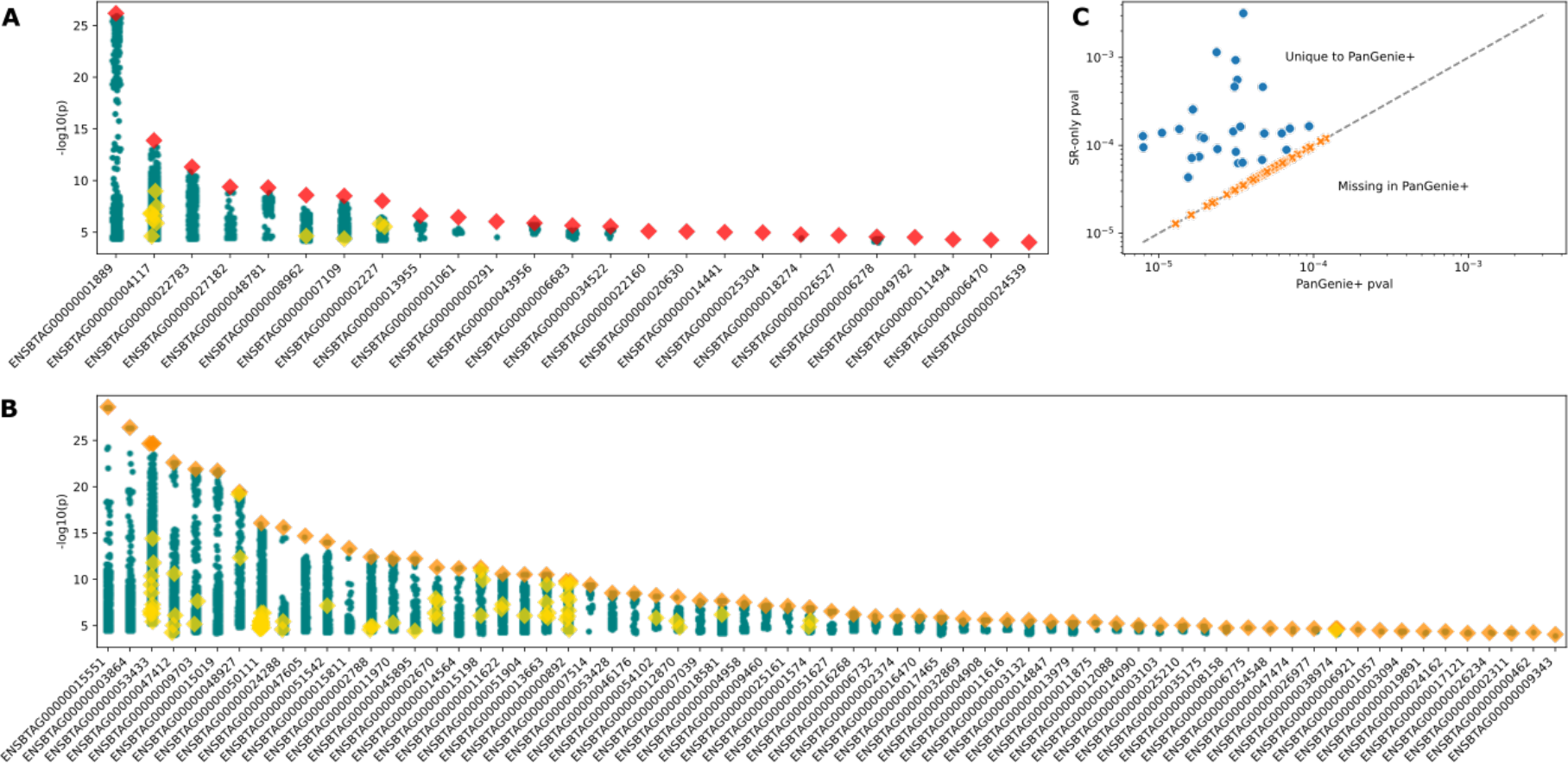
cis-QTL mapping. (A) 25 independent eGene signals with red diamonds denoting SVs as uniquely top hits. Other SVs are shown as yellow diamonds, and small variation are shown as teal circles. (B) 58 independent eGene signals with SVs as top hits in LD with small variants denoted as orange diamonds, with yellow diamonds and teal circles as described in (A). (C) eGenes that are present in only the PanGenie+ dataset or the short read-only DeepVariant dataset. The dashed line indicates equal significance thresholds between the two conditional analyses.

**Figure 4.**
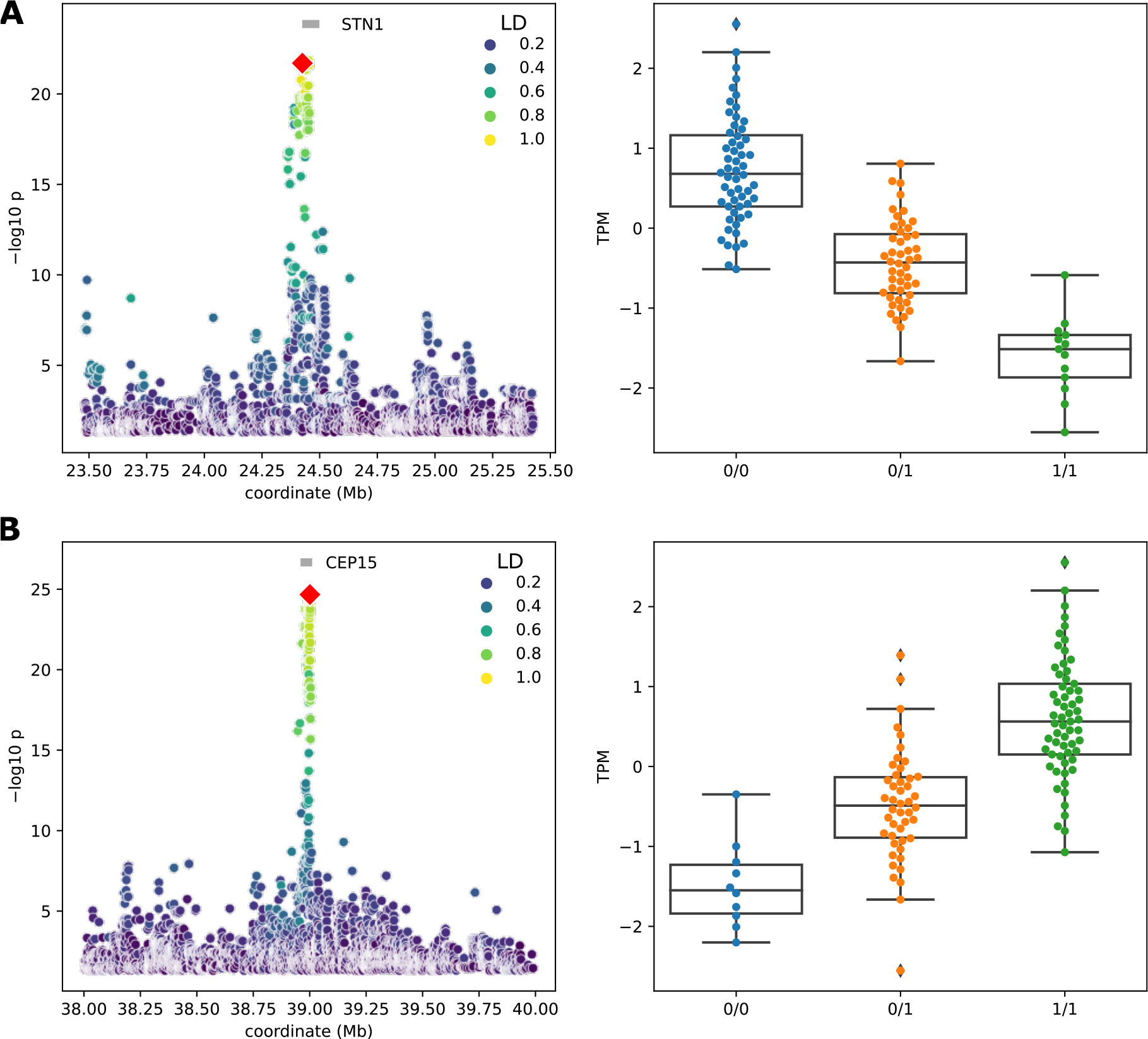
Nominal eQTL association significance (left) and normalized TPM values for the expressed gene (right) for (A) STN1 and (B) CEP15. The red symbol represents the top-associated SV and. Linkage disequilibrium (LD) between the SV and all other variants within the cis-window is indicated with the color gradient.

A 118 bp deletion was strongly associated with *CEP15* (*ENSBTAG00000001889*, encoding centrosomal protein 15) mRNA abundance (Figure 4B). The deleted sequence is a short interspersed nuclear element (SINE). The SV-eQTL was located 8 Kb downstream of the transcription start site of *CEP15* and was 1.4x as significant (p= 6.60e-27) as the closest SNP. The deletion was associated with increased (ß=1.23) *CEP15* expression.

We also examined two prominent insertion SV-eQTL. The expression of *MYH7* (ENSBTAG00000009703) encoding myosin heavy chain 7 was associated with a 388 bp insertion (p=1.27e-22) consisting almost entirely of LINE sequence. The LINE sequence was inserted 8.3 Kb downstream of *MYH7* and increased mRNA abundance (ß=1.37). The expression of *LOC112443864* (ENSBTAG00000053433) encoding MHC class I polypeptide-related sequence B-like was associated with an 11.6 Kb insertion (p= 2.18e-25) containing 2.3 Kb of SINE, LINE, and ERV-LTR elements 7.4 Kb upstream (ß=1.09) (Supplementary Figure 7). Given its location nearby the bovine leukocyte antigen (BoLA) complex, this SV potentially could contribute to eQTL in immune-related tissues.

We realized that the 11.6 Kb insertion affecting *LOC112443864* expression also highlights difficulties in association testing with large SVs. The original pangenome variant panel constructed from the 16 haplotypes contained three near-identical (>99.9% sequence identity) insertion alleles, differing by only several SNPs. Each allele, when considered individually for molQTL mapping after PanGenie-based genotyping, was below the significance threshold for *LOC112443864*, but curating and merging the alleles before genotyping and conducting the eQTL analysis revealed a highly significant peak (Supplementary Figure 8).

### cis-splicing QTL mapping

We performed a similar analysis for splicing QTL (sQTL), now using intron excision ratios as the phenotypes. We tested for associations in 14,243 genes with 46,417 splicing clusters. With the PanGenie+ variant set, we find 3,613,475 associated variants for 16,893 sQTL (7,064 spliced genes [sGenes] and 10,629 splicing clusters), of which 5,366 were SVs and 1,061 were SVs larger than 1 Kb. Again, we found only 11 additional sGenes compared to using the short read-only dataset, but we did find 73 SV-sQTL with 15 sSVs as the unique-top variant (Figure 5A) and 58 sQTL where the top variant is an SV that is in near-perfect linkage disequilibrium with a small variant (Figure 5B, Supplementary Table 1).

**Figure 5.**
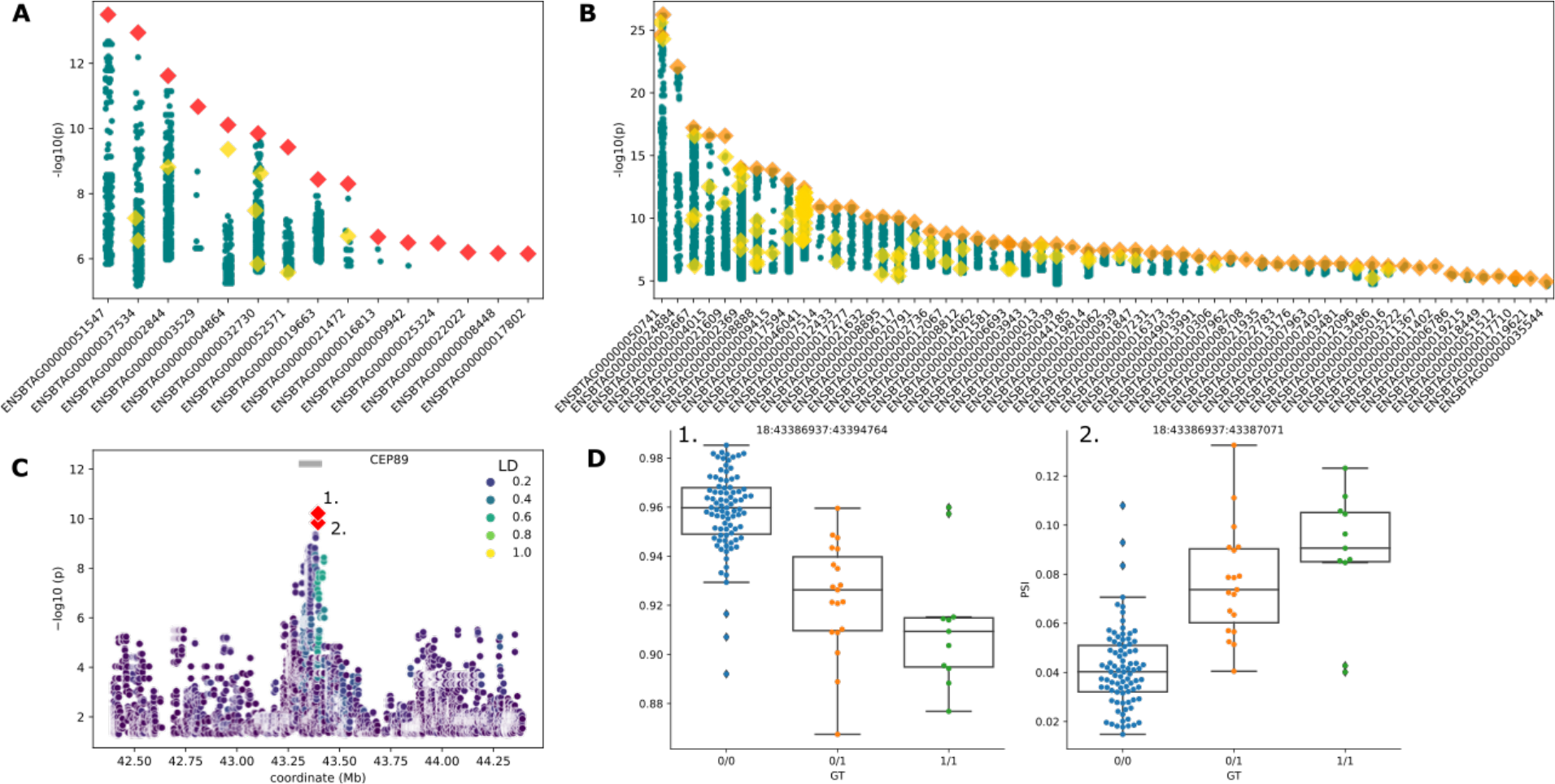
cis-sQTL mapping. (A) 15 independent sGene cluster signals with SVs as the unique-top variant and (B) 58 SVs as top variants in LD with small variants, with the color and marker meanings described in Figure 4. (C) Nominal association significance for CEP89, where the two red diamonds indicate the same variant affecting two separate junction splicings within the sQTL cluster. (D) PSI (percent spliced in) across the two significantly associated junctions (indicated by number from (C)) within the sQTL cluster.

We examined an sQTL for *CEP89* (ENSBTAG00000004864) encoding centrosomal protein 89 in more detail, noting that a 1.3 Kb insertion at Chr18:18:43,395,289 3 Kb downstream of the transcription start site was the top associated variant for a splice cluster containing two splice junctions. This 1.3 Kb insertion was approximately 6 times more significant (p=6.0e-11) than the next highest SNP, with ß=-0.86 (Figure 5C). The inserted sequence was almost entirely an LTR retrotransposon and present in approximately 30% of samples, contributing to alternative splicing (Figure 5D). The two associated splice junctions span the second and third exon of *CEP89* (Supplementary Figure 9). However, the annotation of three *CEP89* transcripts in Ensembl again appears incomplete as Refseq indicates seven *CEP89* isoforms. While the SV does not affect the overall *CEP89* expression (i.e., *CEP89* was not an eGene), it is associated with the abundance of two isoforms suggesting that this sQTL promotes alternative isoform usage and so impacts the relative abundance of distinct CEP89 isoforms.

We also examined an sQTL for *ASAH2* (ENSBTAG00000003529) encoding N-acylsphingosine amidohydrolase 2. The top associated variant was 34 Kb downstream and was a 3.6 Kb insertion and was approximately 150 times more significant (p=1.60e-11) than the next most significant SNP, causing alternative splicing (Supplementary Figure 10) with ß=1.24. The inserted sequence contained nearly 2 Kb of BovB repeats, another transposable element.

The inserted sequence was located within a putative duplication, causing potential misalignments of short reads and thus erroneously calling a C-to-T transition at 26:8694035. This variant has previously been reported (EVA: rs385128608). However, long read alignments more strongly support the 2 Kb insertion detected through the assemblies, although even these were complicated to validate (Supplementary Figure 11). Although the SV genotyped through PanGenie appears to be the most accurate, the limited LD observed with adjacent small variations may also indicate imperfect genotyping due to limited unique *k*-mers in the region. More generally, there are 4,338 instances where a SNP and SV share a starting genomic coordinate, of which 1,851 (42.7%) occur in regions identified as VNTRs (Leonard et al. 2023), where short reads can easily misalign and appear as motif variation rather than insertions/deletions of additional repeats. There are 599 and 425 SV-e/sVariants respectively overlapped by SNPs, of which 61 and 85 respectively are highly significant (p<1e-10).

## Discussion

Including structural variation for expression and splicing QTL mapping provides additional insight into functional genomic elements over just including the small variation that can confidently be called from short reads. PanGenie is one approach that enables effective and accurate genotyping of small and structural variation, discovered through a relatively small set of assembly haplotypes, into a much larger short read cohort. Even 8 individuals (16 haplotypes) appear sufficient to capture 69% of SVs in 25 unrelated samples, with most of the missing SVs occurring at low frequency in the 25 samples. The assemblies also could detect larger insertions compared to using the samples’ raw sequencing directly with read-based approaches, as recently observed (Harvey et al. 2023). Furthermore, we identify cases like an SV-sQTL for *ASAH2* where SNPs (including some reported in public databases) may actually be SVs. PanGenie can resolve such cis-e/sQTL associations with the genotyped SVs correctly called from high quality assemblies. Given a larger number of initial assemblies, association mapping may further improve by only using PanGenie genotyped variation and not supplementing with occasionally erroneous short read-called variation.

We observe extensive LD between SNPs and SVs, as reported elsewhere (Lee et al. 2023; Zhou et al. 2022b), which limits the number of e/sGenes that were uniquely discovered by the PanGenie variant set. However, including SVs revealed equally or more significant top associations, and in some cases were more compelling causal candidates (e.g., deletion of an entire exon) than the tagging SNPs. This is particularly true for 53 and 41 insertion SV-e/sQTL respectively, which can only be interrogated through long reads or assembly-based approaches. Still, the variant set lacks a moderate fraction of SVs segregating in this cattle population, particularly rare alleles that might have an especially strong impact on gene expression and splicing (Wagner et al. 2023; Li et al. 2017a), and so a full long read cohort may provide greater power to find untagged SV-QTLs.

Several of the SV-QTL examined in detail (e.g., *STN1*, *CEP89*, and *ASAH2*) contain inserted or deleted sequences that are largely comprised of transposable elements, like ERV-LTR, BovB, and hobo transposons. More generally, we found 51 out of 92 (55.4%) SV-eQTL and 37 out of 73 (50.7%) SV-sQTL contained transposable elements (Supplementary Table 2). While many of these e/sQTL were also associated with SNPs that were in LD with SVs, transposable elements are widely reported to be able to mediate expression (Elbarbary et al. 2016; Kelly et al. 2022; Almeida et al. 2007; Platt et al. 2018), and so are strong candidates for being the causal variants. Given the SV size distribution spike around the size of LTR elements, it is likely such transposable elements will increasingly be identified as a driving force behind bovine phenotypic diversity.

Association mapping with SVs is not just a simple extension to using SNPs, due to SVs’ greater proclivity of having highly similar but distinct alleles. Larger SVs (e.g., >1 Kb) are likely to appear multiallelic across older assemblies or individual high-quality reads (typically quality value of ∼30 or 1 error expected per 1 Kb). Distinguishing technical noise (errors in reads/assemblies) from meaningless biological variation (differences in allele have no functional consequence) from meaningful biological variation (differences in allele may functionally impact gene regulation) is an open and challenging question. Addressing this question is particularly critical for pangenomes containing diverse (sub)species, as multiallelic but similar SVs become increasingly common, which can dilute significant associations below their thresholds.

We also confirm recent results that at moderate coverages (approximately 10-fold), HiFi reads can replace short reads for small variant calling, while are also able to accurately call SVs (Kucuk et al. 2023; Harvey et al. 2023). The former is especially true in highly repetitive regions like centromeric satellites or tandem repeats, which have largely been challenging to assess with short reads even using dedicated tools. As such, future large efforts, like the cattle Long Read Consortium (cattleLRC, (Nguyen et al. 2023)), will likely be able to assess nearly all genomic variation from only a single data source of accurate long reads. However, in the intermediate future while cohort-scale costs are prohibitive, we demonstrate that several assemblies and PanGenie can greatly improve the quality and accuracy of e/sQTL and related analyses.

## Methods

### HiFi sequencing

We extracted high-molecular weight DNA from blood of two animals with the Qiagen MagAttract HMW kit, following manufacturers protocols. PacBio HiFi libraries were generated and sequenced on three SMRT cells each by the Functional Genomic Center Zurich (FGCZ).

Testis tissue from 25 additional BSW/OBV individuals was sampled from a commercial abattoir in Zürich, Switzerland. We extracted high-molecular weight DNA with the Monarch HMW Extraction Kit for tissue (New England BioLabs) and followed manufacturer recommendations. DNA fragment length and quality was assessed by the FGCZ with the Femto Pulse System (Agilent). PacBio HiFi libraries were produced and sequenced on one SMRT cell per individual with a Sequel IIe.

### Genome assembly

Four Original Braunvieh and four Brown Swiss haplotypes were assembled from publicly available data (project code PRJEB42335). In addition, we assembled four Brown Swiss haplotypes from new HiFi data (accession codes ERS15606279 and ERS15606280) from two F1s. We used hifiasm (v0.19.4-r575) (Cheng et al. 2021) to generate the haplotype-resolved assemblies, using default parameters and providing parental *k*-mers of size 31 counted by yak (v0.1-r66-dirty) for the two trios. We scaffolded the resulting contigs to ARS-UCD1.2 using RagTag (v2.1.0) (Alonge et al. 2022) with the additional parameters “-cx asm20”.

### PanGenie genotyping

We created the variant panel from the 16 cattle assemblies following the approach laid out by PanGenie. Briefly, we aligned each assembly to ARS-UCD1.2 with minimap2 (v2.24-r1122) (Li 2018) with parameters “-ax asm20-m 10000-z 10000,50-r 50000-end-bonus=100 −O 5,56-E 4,1-B 5”, followed by calling haploid variants for each haplotype with paftools.js. Variants were merged into diploid calls and filtered according to PanGenie. We additionally modified the merging step to consider SVs with >98% sequence identity to be part of the same cluster and take the first SV of the cluster as the allele.

We genotyped 307 short read samples using PanGenie (v.2.1.1) (Ebler et al. 2022) with the pangenome variant panel using default parameters. Each VCF was then merged using bcftools (v1.17) merge (Danecek et al. 2021).

### Small variant calling

We aligned short read samples to ARS-UCD1.2 using bwa-mem (v0.717) (Li 2013) using the -M flag, followed by coordinate sorting and de-duplicating with samtools (v1.17) (Danecek et al. 2021). Variants were called per-sample using DeepVariant (v1.5.0) (Poplin et al. 2018) with the “WGS” model and jointly genotyped and filtered using GLnexus (v1.4.1) (Yun et al. 2021) with the “DeepVariantWGS” configuration. Sporadically missing variants were then imputed using Beagle (v5.4) (Browning et al. 2018).

We aligned long read samples to ARS-UCD1.2 using minimap2 with the parameters “-ax map-hifi” and converted to BAM files as described above. We called variants as above, except using the “PACBIO” model for DeepVariant.

### Structural variant calling

We called and jointly genotyped SVs using Sniffles (v2.0.7) (Smolka et al. 2022) on the aligned long read files with the parameter “--min_sv_len=50”. For the assembly haplotype samples, we additionally used the “--phased” parameter. We filtered out BND-type variants as well as variants exceeding 100 Kb with bcftools view.

### Variant analyses

We used bcftools to merge the DeepVariant short read-called variants with the PanGenie genotyped variants for the 307 samples, using the concat command with the “-D” flag to remove duplicate variants (giving allele/genotype priority to DeepVariant). Indels were left-normalised with bcftools norm.

We assessed genotype accuracy using hap.py (v0.3.15) (https://github.com/Illumina/hap.py), using the short read-called variants as truth and the HiFi-called variants as query. We determined the overlap of the two variant sets using bcftools isec -c some -n +1 to allow partial overlapping of multiallelic sites, followed by determining the proportion in centromeric satellites using bedtools (v2.30.0) (Quinlan and Hall 2010) intersect on those positions and annotated regions. We determined if multiples SVs were “the same” using Jasmine (v1.1.5) (Kirsche et al. 2021), allowing intersections up to the smaller of max_dist_linear=1 (proportional to SV size) and max_dist=1000 (1 Kb).

We used bcftools mendelian2 plugin to assess mendelian inconsistency rates.

### RNA sequencing and alignment

RNA from 117 testis samples were sequenced from paired-end total RNA libraries, as described in (Mapel et al. Companion paper). Briefly, the sequencing reads were trimmed using fastp, and aligned to ARS_UCD1.2 and the Ensembl gene annotation (release 104) with STAR (version 2.7.9a) (Dobin et al. 2013). We produced an additional set of alignments with the flag -waspOutputMode to account for allelic mapping bias for sQTL analyses.

### QTL analyses

Gene quantification and covariate files were processed for e/sQTL analyses as described in (Mapel et al. Companion paper). Briefly, to quantify gene-level expression in TPM we used QTLtools quan (Delaneau et al. 2017) and to infer gene-level read counts we used featureCounts (Liao et al. 2014). We removed lowly expressed genes and only included genes with ≥ 0.1 TPM in ≥20% of samples and ≥ 6 reads in ≥ 20% of samples. Filtered expression values were quantile normalized and inverse normal transformed for downstream analyses.

For splicing quantification, we considered intron-excision values from intron clusters identified. Specifically, we identified exon-exon junctions from WASP-filtered reads with Regtools (Cotto et al. 2023), followed by using Leafcutter (Li et al. 2017b) to construct intron clusters and an altered “map_clusters_to_genes.R” script to map clusters to the cattle gene annotation (Ensembl release 104). We filtered introns with read counts in <50% of samples, introns with low variability across samples, and introns with fewer than max(10, 0.1*n*) unique values (where *n* is sample size). We used the “prepare_phenotype_table.py” script from Leafcutter to normalize filtered counts and produce files for sQTL mapping.

We filtered variants with MAF < 0.01 and split multiallelic sites using bcftools view and norm respectively. We performed all association testing (for both e/sQTL) using QTLtools (v1.3.1) (Delaneau et al. 2017). Permutation analyses were performed using a 1 Mb cis-window 2000 times with a false discovery rate of 0.05, which determined the significance thresholds for each gene in the conditional pass. Nominal association was performed using a significance threshold of 0.05. LD scores for specific variants were calculated using plink v1.9 (Chang et al. 2015).

The abundance of Refseq (version 106, GCF_002263795.1) transcripts was quantified using kallisto (version 0.46.1 (Bray et al. 2016)), and aggregated to the gene level using the R package tximport (Soneson et al. 2016).

### Data Access

Short DNA and RNA sequencing reads are available in the ENA database at the study accessions PRJEB28191 and PRJEB46995. The HiFi data generated in this study are available at the study accessions PRJEB46995 and PRJEB42335. Parental short reads of two F1s (SAMEA8565097, SAMEA32982418) are available at the study accession PRJEB18113 under sample accessions SAMEA8565028 and SAMEA8565098 for the first and SAMEA32980918 and SAMEA32981668 for the second F1. The HiFi reads of the F1s are available under accessions ERS15606279 and ERS15606280. All scripts are available online (https://github.com/AnimalGenomicsETH/pangenome_molQTL).

## Acknowledgements

We thank Dr. Anna Bratus-Neuenschwander and Dr. Catharine Aquino from the ETH Zurich technology platform FGCZ (https://fgcz.ch) for DNA fragment analysis and DNA and RNA sequencing. We also thank Eirini Lampraki from Pacific Biosciences for DNA sequencing. We thank Dr. Cord Drögemüller (University of Bern) for providing blood samples.

## Author contributions

A.S.L. and H.P. conceived the study. X.M.M performed the DNA and RNA sequencing. A.S.L. constructed the genome assemblies and created the small, structural, and PanGenie variant sets. A.S.L. ran the e/sQTL association analyses with input from X.M.M. A.S.L. and H.P. performed detailed analyses on specific QTL. A.S.L. and H.P. wrote the manuscript. All authors read and approved the final manuscript.

