## Supplementary material for "Pangenome genotyped structural variation improves molecular phenotype mapping in cattle": Supplementary_material.docx

Supplementary Figure 1. Series of 4 IGV screenshots for one sample with haplotype-binned HiFi reads (upper panel) and haplotype-resolved assemblies (lower panels). (a-b) Two examples where haplotype-resolved assemblies can confidently identify an SV but HiFi read-based SV calling fails, and (c-d) two examples where read-based calling matches the SVs called by assemblies.

Supplementary Figure 2. PCA for small (A) and structural (B) variants genotyped in the 307 samples. Slightly more variance was explained by the top 10 PCs for small variants compared to structural variants. Breed structure is revealed for Brown Swiss and Original Braunvieh, with crosses between these breeds (Braunvieh) or crosses with a different breed (Cross) distributed between the two primary clusters.

Supplementary Figure 3. New unique SVs per additional sample added to the cohort (black markers). The grey fitted line is taken from Figure 2b. The red and orange horizontal lines respectively indicate expected discovery of 1000 and 100 new unique SVs per sample, intersecting the fitted line at approximately 14 and 108 samples.

Supplementary Figure 4. (A) Both HiFi and short reads (SR) alignments confidently span almost the entirety of the autosomes, while HiFi performs better for the typically more complex and repetitive sex chromosomes (X and Y). (B) HiFi-based alignments called more variants than short read-based alignments at approximately 10-fold coverage, particularly in the sex chromosomes.

Supplementary Figure 5. Allele frequency distribution for SNPs, Indels, and SVs in the PanGenie+ variant set, across the 307 samples.

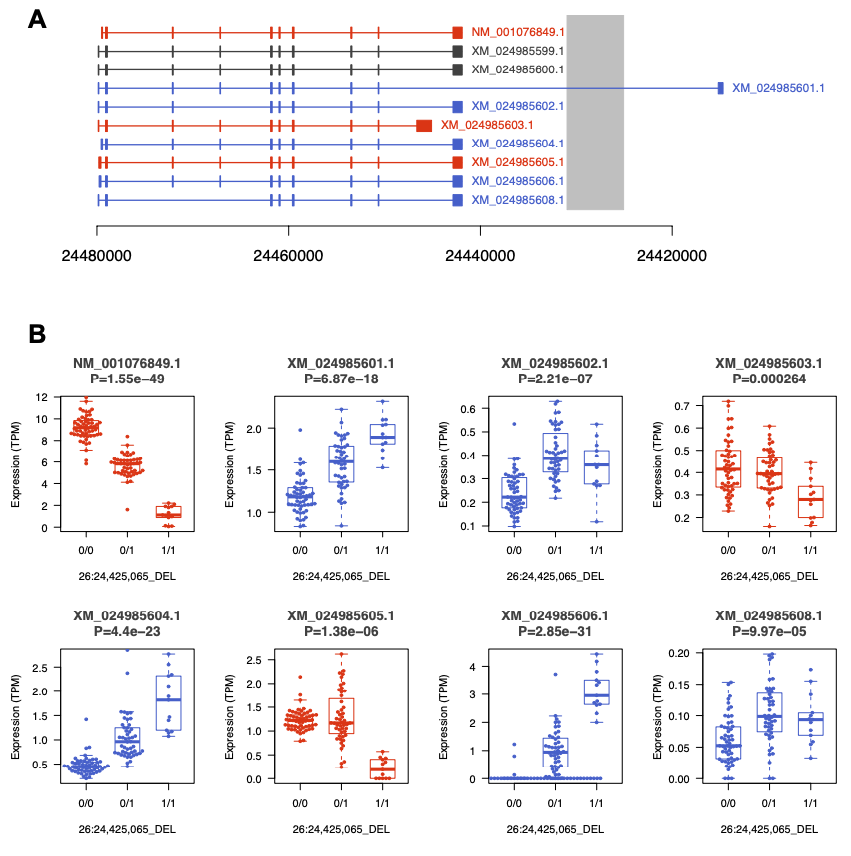

Supplementary Figure 6. A 5.9 kb deletion is associated with STN1 expression. (A) Structure of ten STN1 isoforms that are annotated in Refseq (version 106). The grey box indicates the position of the deletion. Boxes represent exons. Red and blue color indicates transcripts whose expression is respectively reduced and increased by the deletion. Transcripts that are not expressed are grey-colored. (B) Impact of the deletion on the expression of eight STN1 isoforms. Red and blue color indicates transcripts whose expression is respectively reduced and increased by the deletion. The P values are from a linear regression of TPM values on the genotype (coded as 0, 1, 2).

Supplementary Figure 7. Nominal eQTL association significance (left) and normalized TPM values for the expressed gene (right) for (A) MYH7 and (B) LOC112443864.

Supplementary Figure 8. (A) Bandage plot for an 11.6 Kb insertion present in seven out of eight samples. There were three distinct alleles, but with >99.9% sequence identify, differing only by six SNPs and one 2 bp indel across the entire SV. (B) Nominal significance of association for the SV and ENSBTAG00000053433. Given the high sequence identity and limited number of unique k-mers, PanGenie could not consistently genotype the 3 different alleles, and so mean and standard deviation are calculated for 5 replicates. The dashed line indicates the conditional significance threshold calculated from the PanGenie+ dataset. Only the merged PanGenie variant was significant, while the three near-identical alleles were all insignificant due to association dilution.

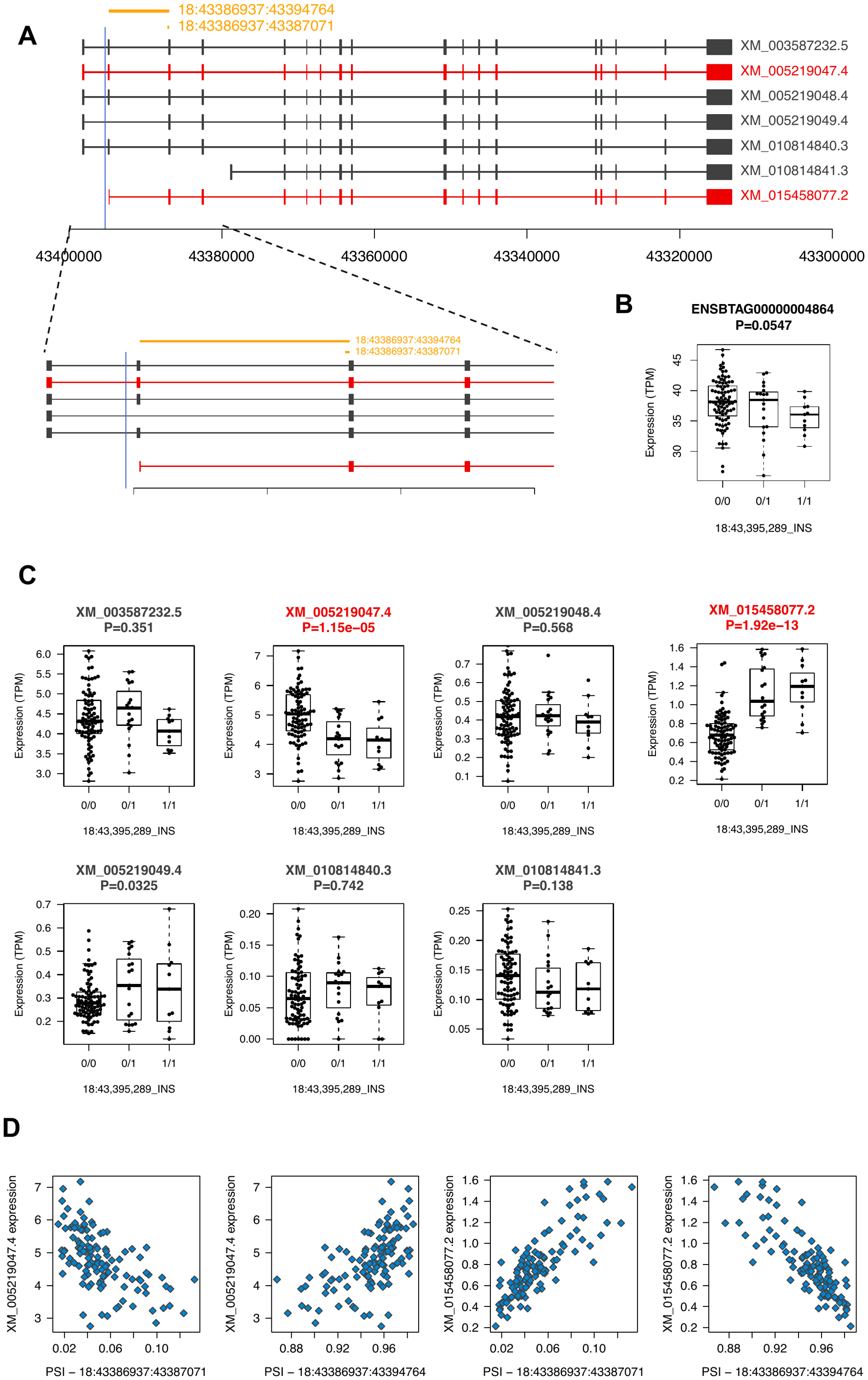

Supplementary Figure 9. An intronic insertion is associated with CEP89 splicing and transcript-level expression. A) Structure of seven CEP89 isoforms that are annotated in Refseq (version 106). The blue vertical line indicates the position of the insertion. Boxes represent exons. Red color indicates two transcripts whose expression is impacted by the insertion. The orange lines represent the two splice junctions that are associated with the insertion. B) Gene-level expression (quantified in TPM) of CEP89 (ENSBTAG00000004864) is not affected by the insertion. The P value is from a linear regression of TPM values on the genotype (coded as 0, 1, 2). C) Impact of the insertion on the expression of seven CEP89 isoforms. Red color indicates two isoforms (XM_005219047.4, XM_015458077.2) whose expression is significantly (P<0.05/7) associated with the insertion. The P values are from a linear regression of TPM values on the genotype (coded as 0, 1, 2). D) Correlation between the expression of the two significant transcripts and the percent-spliced-in (PSI)-values of the two splice junctions.

Supplementary Figure 10. (A) Nominal association significance for ASAH2, where the two red diamonds indicate the same variant affecting two separate junction splicings within the sQTL cluster. (B) PSI (percent spliced in) across the two significantly associated junctions (indicated by number from (A)) within the sQTL cluster.

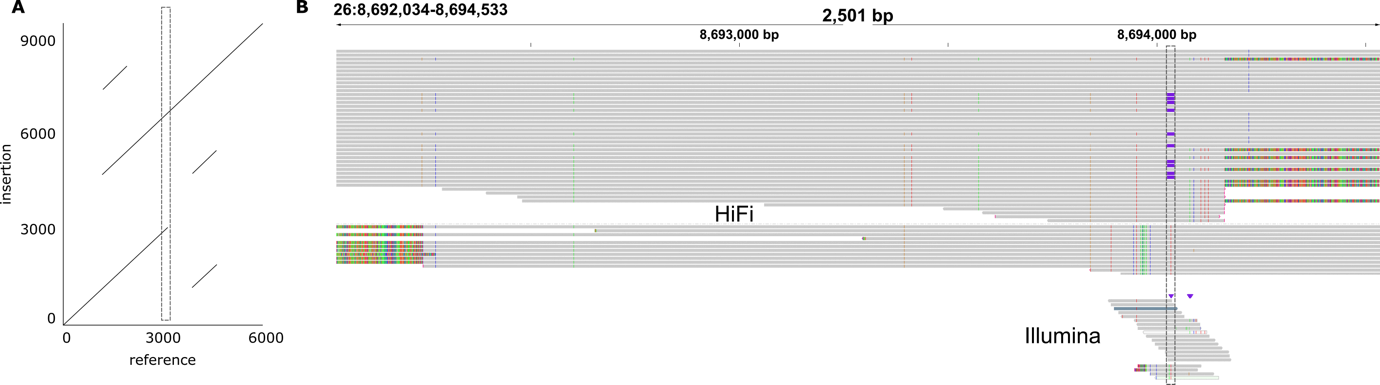

Supplementary Figure 11. a) Dotplot of 6 Kb of the reference sequence (26:8691035-8697035) against the syntenic region containing the 3.6 Kb insertion, where the dotted box indicates the start of the SV. The additional off-diagonal elements indicate duplications. b) IGV screenshot for HiFi (top) and Illumina (bottom) sequencing for the same individual who was heterozygous for the SV, where the dotted box again indicates the start of the SV. Within the dotted box, purple rectangles indicate the insertion SV allele, and red lines indicate the erroneous C-to-T SNP. All HiFi reads not containing the insertion have clear rainbow-banding patterns at their ends, indicating soft-clipping and lower quality alignment compared to reads clearly showing the insertion. Illumina reads were generally too short to confidently state the SNP-reads were misaligned, and so the SNP looks valid.

Supplementary Table 1. SV-QTL candidates for expression and splicing phenotypes, with conditional p-values and effect sizes.

External file

Supplementary Table 2. Percentages of repeat elements across all ARS-UCD1.2 autosomes (Genome-wide) or the top-associated SV-e/SQTL. Statistical significance was calculated using a one-sided Fisher’s exact test on the total number of bases annotated as the respective elements between genome-wide and eQTL and genome-wide and sQTL. Significant p-values are indicated by *.

|  | DNA transposons | SINE | LINE | LTR | nonLTR-RTE | Total |
| --- | --- | --- | --- | --- | --- | --- |
| Genome-wide | 1.23 | 8.11 | 15.08 | 3.05 | 4.94 | 40.40 |
| eQTL | 1.36* | 6.47 | 34.44* | 3.07 | 6.13* | 59.40* |
| sQTL | 0.55 | 5.83 | 39.57* | 17.65* | 5.83* | 78.26* |
