## Supplementary figures and images for "Pangenome genotyped structural variation improves molecular phenotype mapping in cattle"

### Supplementary_Fig_S1.pdf

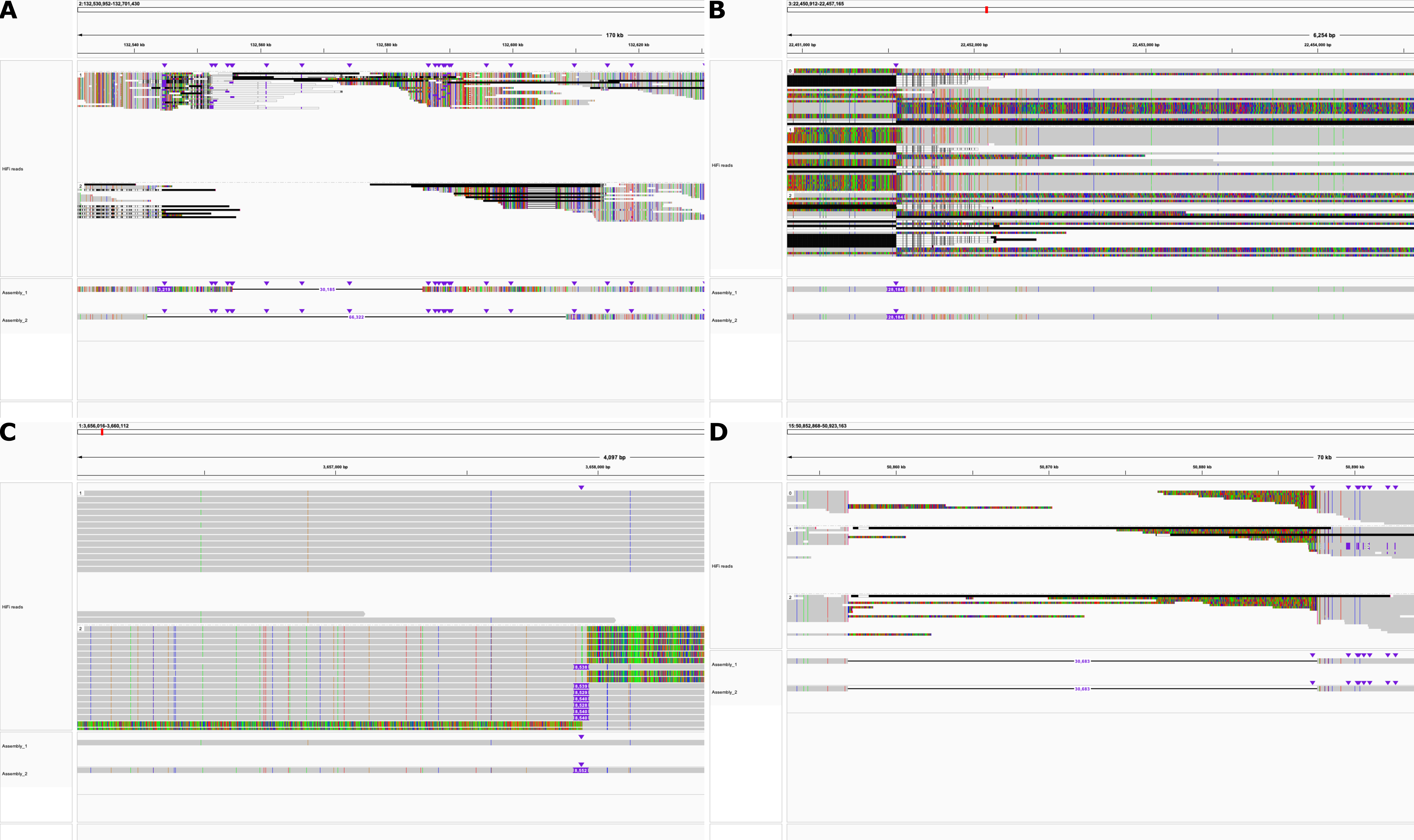

### Supplementary_Fig_S2.pdf

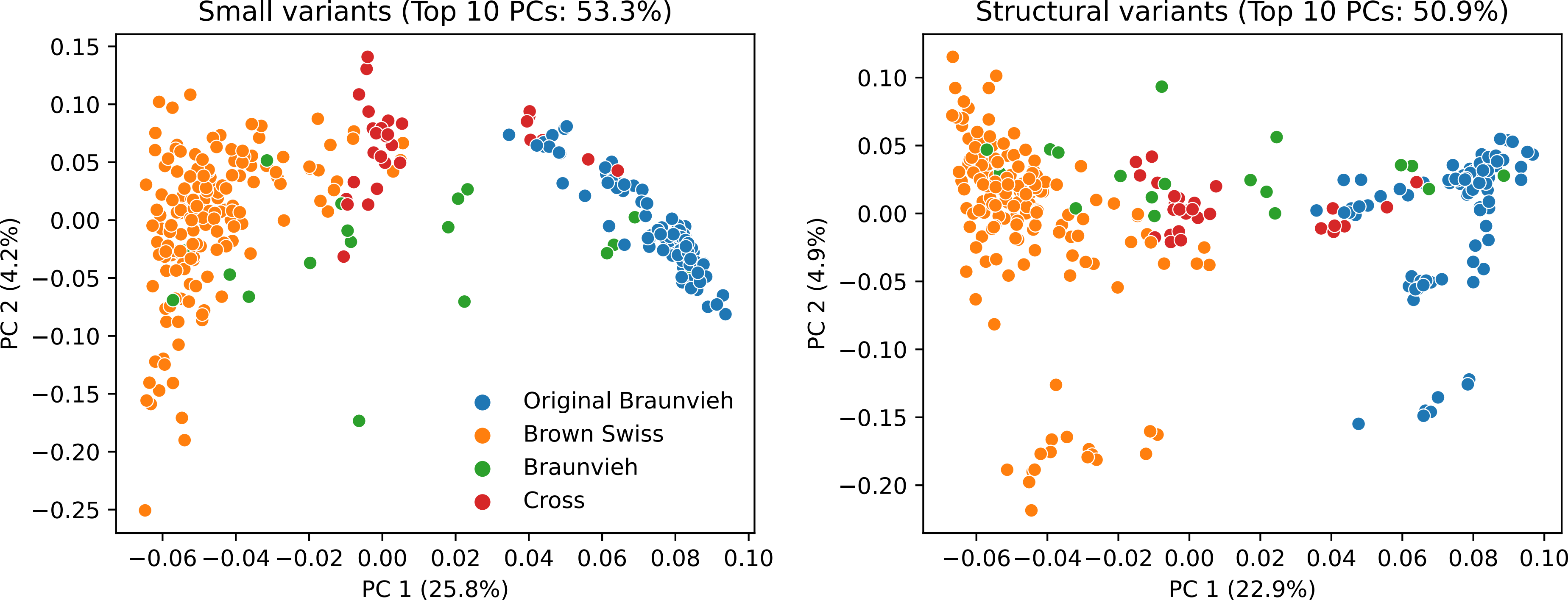

### Supplementary_Fig_S3.pdf

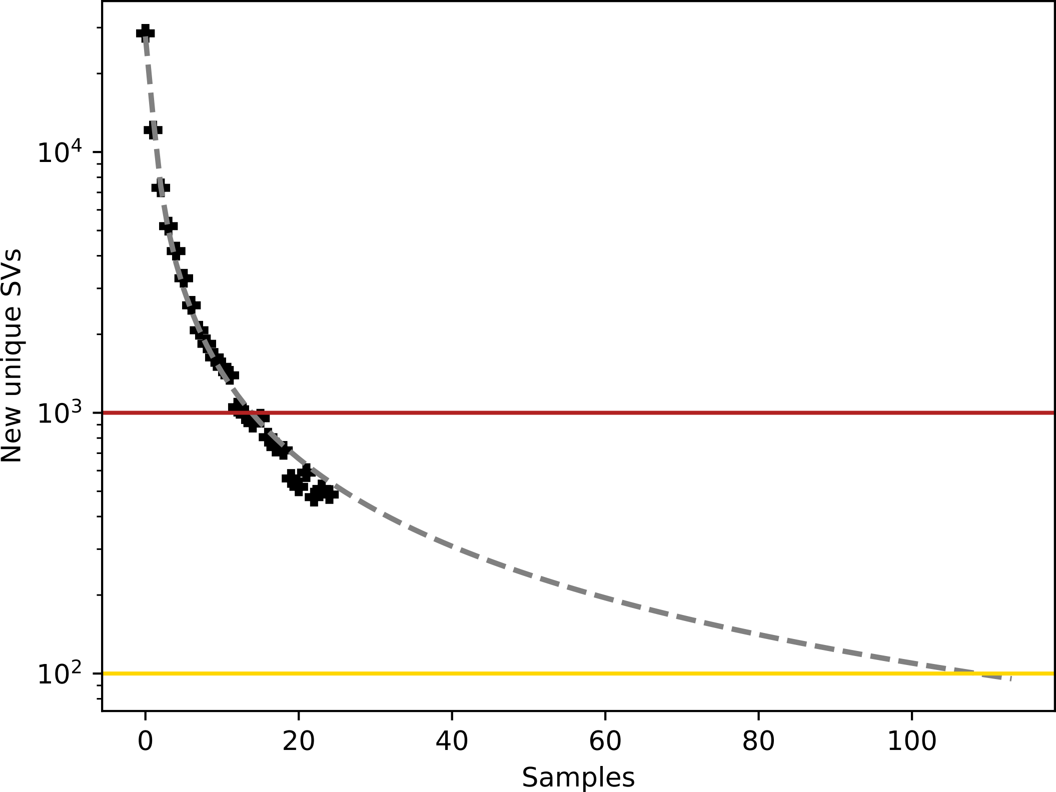

### Supplementary_Fig_S4.pdf

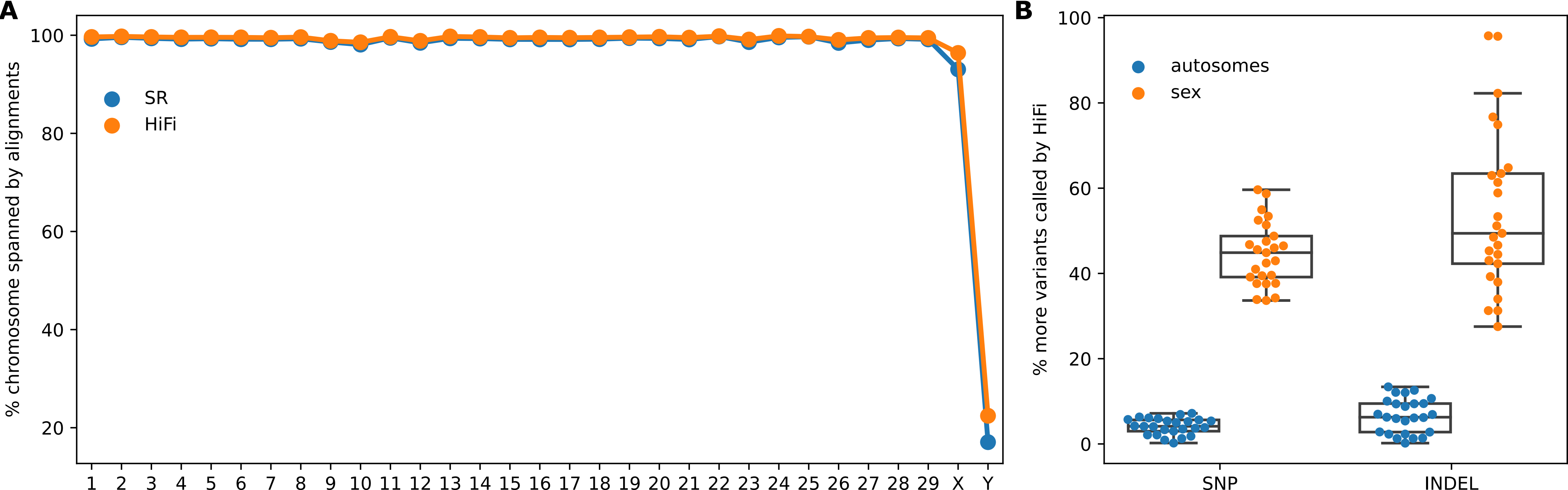

### Supplementary_Fig_S5.pdf

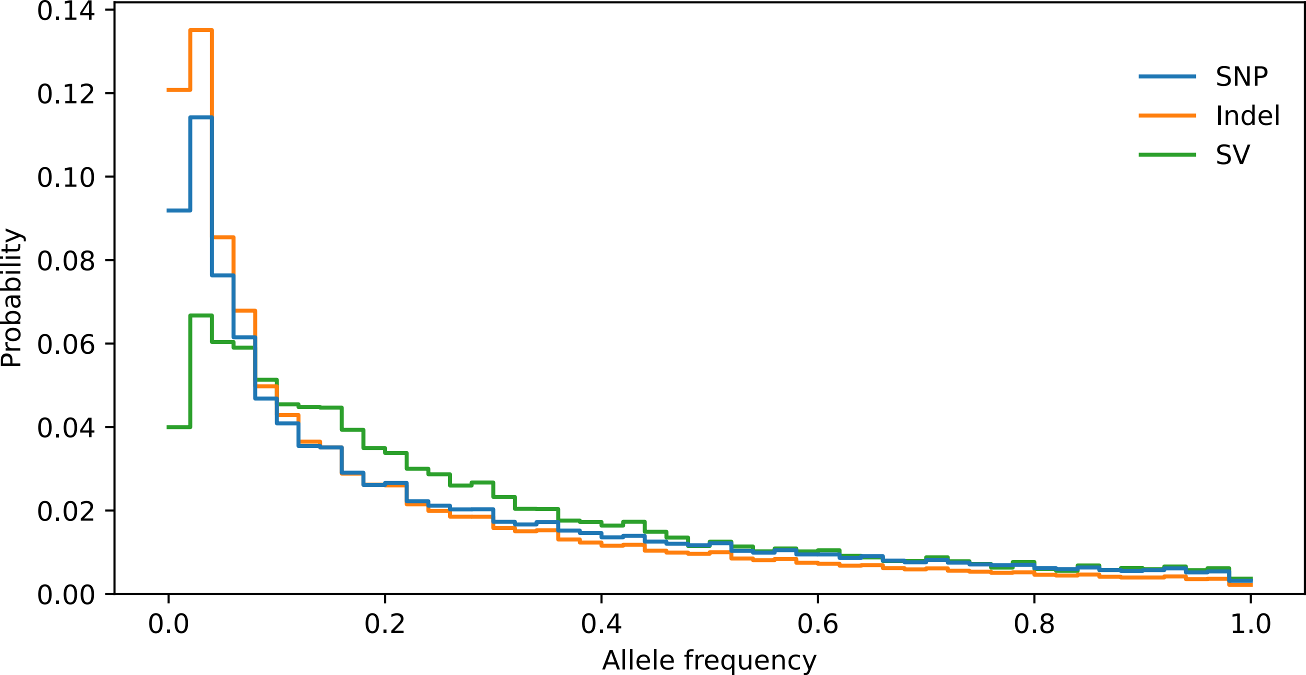

### Supplementary_Fig_S6.pdf

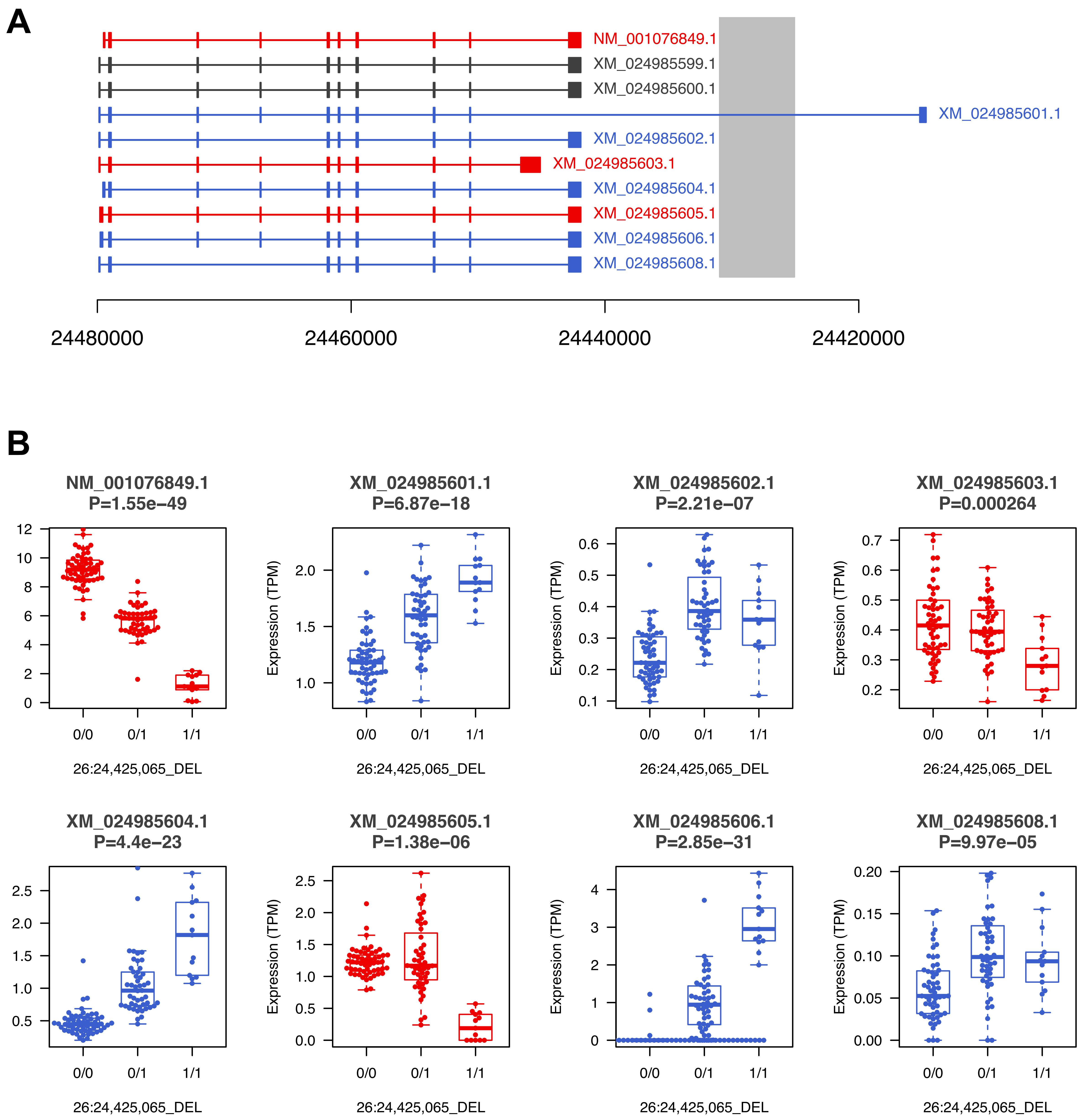

### Supplementary_Fig_S7.pdf

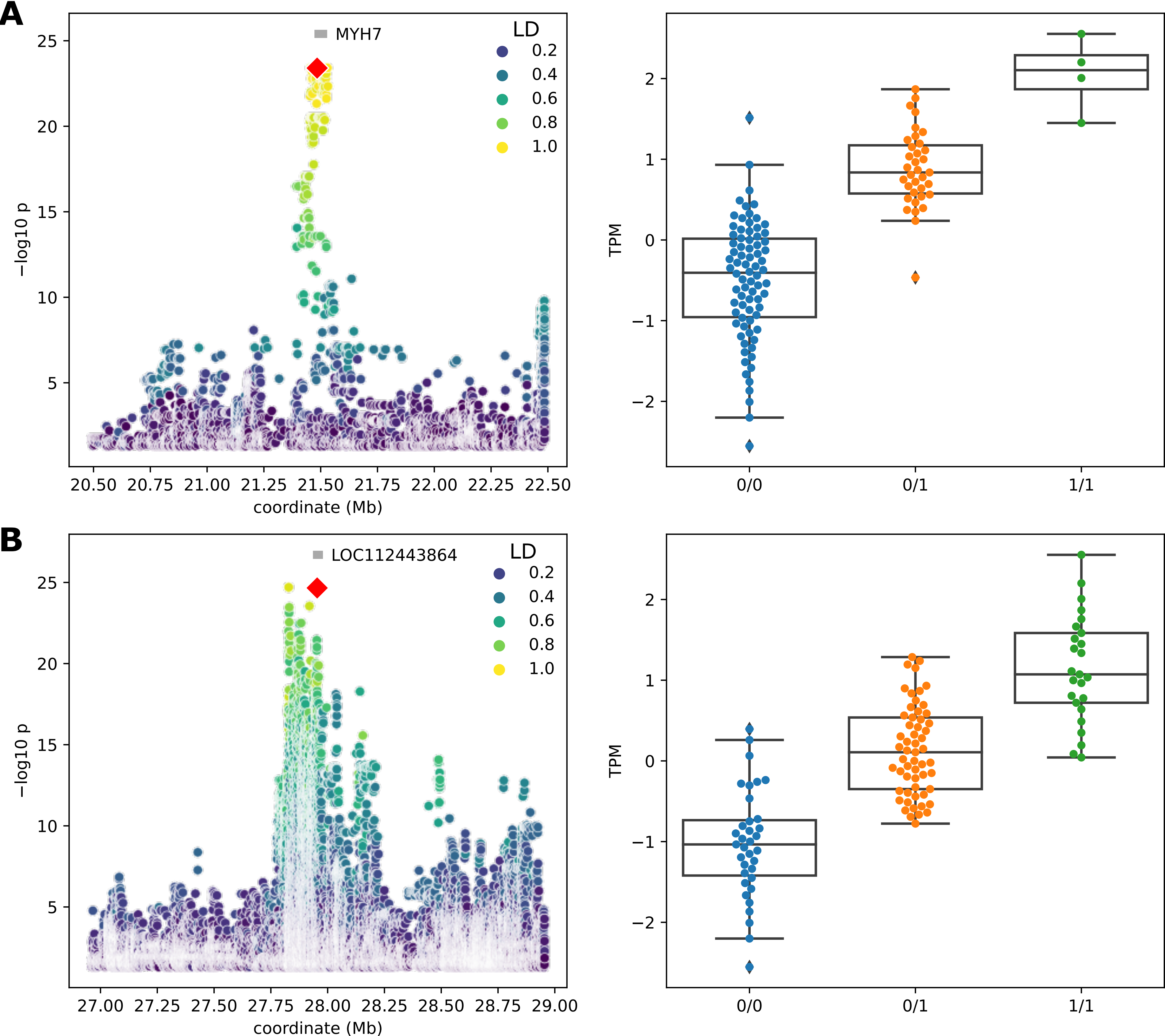

### Supplementary_Fig_S8.pdf

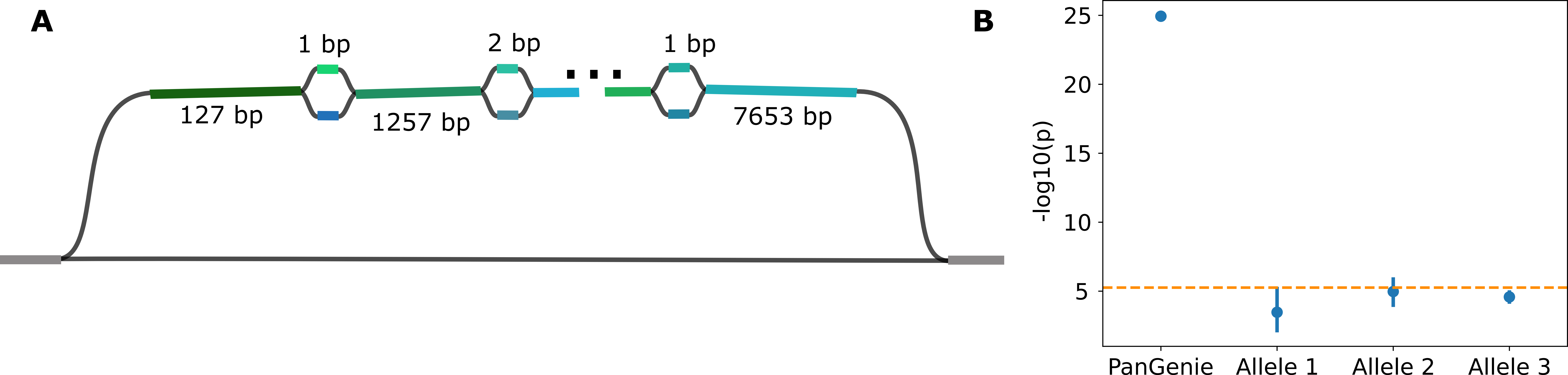

### Supplementary_Fig_S9.pdf

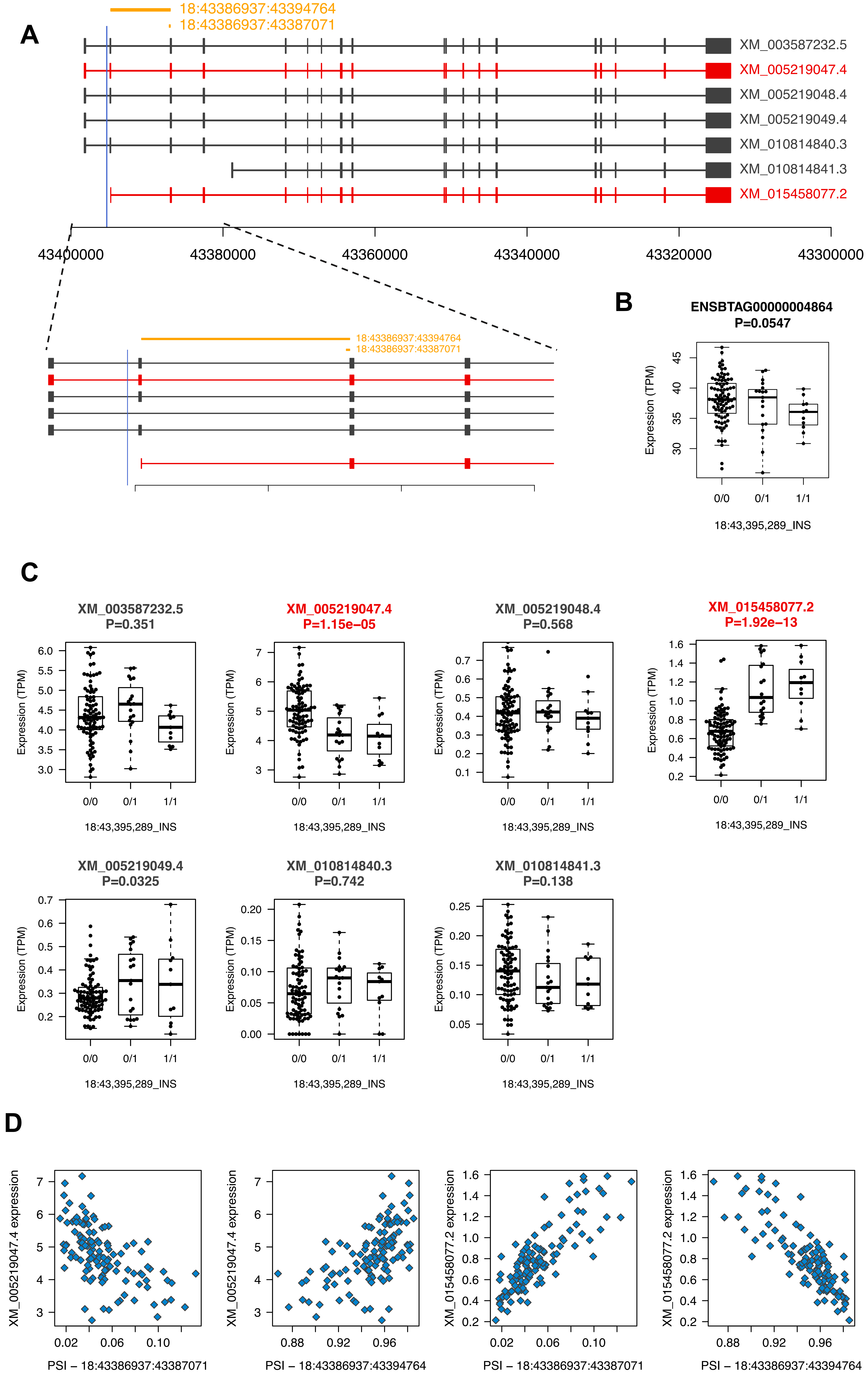

### Supplementary_Fig_S10.pdf

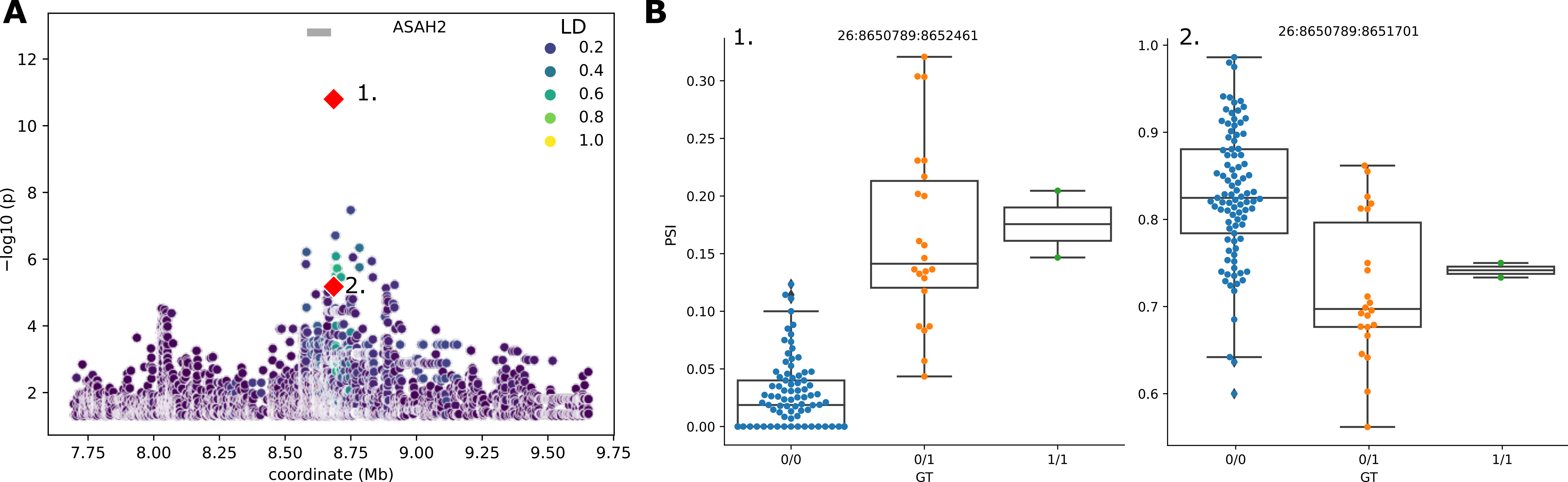

### Supplementary_Fig_S11.png

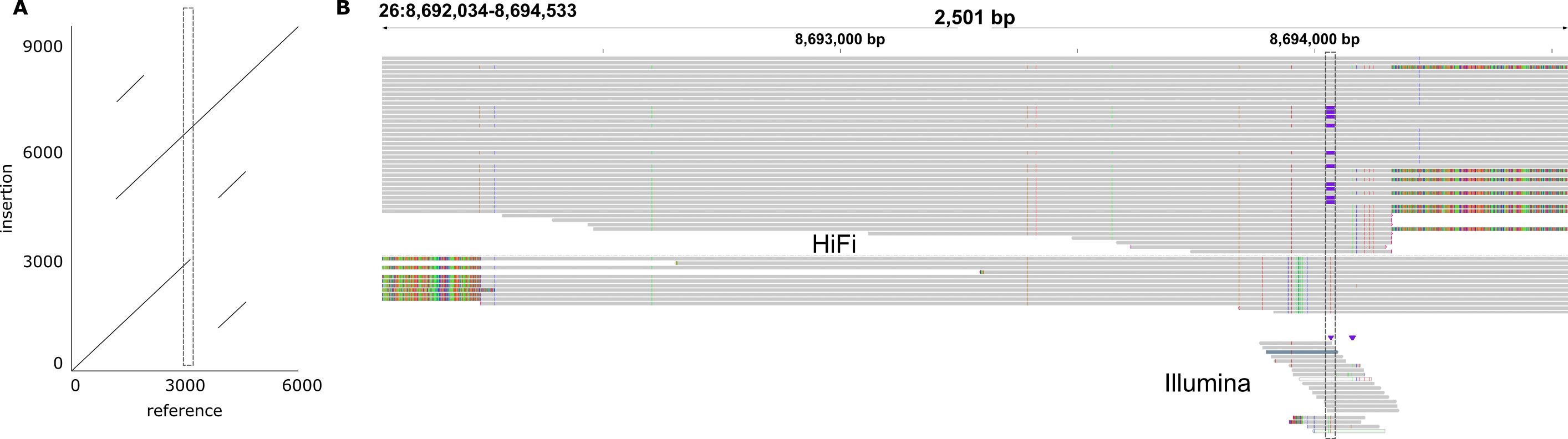
